# Mechanisms of biodiversity between *Campylobacter* sequence types in a flock of broiler-breeder chickens

**DOI:** 10.1101/2021.04.14.439797

**Authors:** Thomas Rawson, Frances M. Colles, J. Christopher D. Terry, Michael B. Bonsall

## Abstract

A long-term study of *Campylobacter* sequence types was used to investigate the competitive framework of the *Campylobacter* metacommunity, and understand how multiple sequence types simultaneously co-occur in a flock of chickens. A combination of matrix and patch-occupancy models were used to estimate parameters describing the competition, transmission, and mortality of each sequence type. It was found that *Campylobacter* sequence types form a strong hierarchical framework within a flock of chickens, and occupied a broad spectrum of transmission-mortality trade-offs. Upon further investigation of how biodiversity is thus maintained within the flock, it was found that the demographic capabilities of *Campylobacter*, such as mortality and transmission, could not explain the broad biodiversity of sequence types seen, suggesting that external factors such as host-bird health and seasonality are important elements in maintaining biodiversity of *Campylobacter* sequence types.

## Introduction

*Campylobacter* are one of the most frequent causes of food poisoning in the UK^1,2^, presenting an estimated £50 million direct economic burden to the UK^3^. The most commonly identified route of transmission to humans is via poultry meat^4^, with seventy three percent of UK supermarket chicken carcasses shown to carry the bacteria^5^. Whereas some foodborne pathogens, such as *Salmonella*, have been shown to proliferate primarily at the slaughterhouse^6^, *Campylobacter* instead emerge and spread rapidly at the farm level^7,8^. As a result, limiting the spread of *Campylobacter* within poultry farms has been one of the primary goals of the Food Standards Agency (FSA) across the last ten years^9^, where attempts to-date have focused on biosecurity measures^10,11^, such as employing anti-bacterial ‘boot dips’ at the entrance to chicken houses, and greater stress placed on farmers to practise consistent hand-washing and facility cleaning. Since *Campylobacter* have been shown to spread from a single bird, to an entire flock, in as little as one week^12^, the thinking behind such prevention methods is to minimise the chance of the bacteria entering the flock in the first instance. Such measures have proved largely ineffective^13–15^, prompting calls for greater study into the ecology of this microbe^11,16^, in the hope of gaining insight into how it can be controlled.

Different strains of *Campylobacter* are commonly categorised by sequence type (ST); genotyping samples by multi-locus sequence typing (MLST) of seven house-keeping genes^17,18^. Broiler flocks (birds grown for their meat) are grown for only a short time, ranging from roughly five weeks for standard flocks, to 12 weeks for organic flocks^19^. Yet despite this short window of time available for *Campylobacter* to colonise a flock, multiple STs are commonly observed simultaneously within a broiler flock^20–22^. For multiple STs to co-occur within a flock for several weeks implies the presence of regulatory mechanisms driving the sustained biodiversity within the flock, that have not yet been identified, let alone studied in depth.

Understanding the inter-strain competition mechanisms amongst different strains of *Campylobacter* can both aid understanding of the host-pathogen relationship, but also presents new opportunities in disease control. Understanding how certain STs may be excluded from colonising a flock by pre-established STs creates the opportunity for manipulation of these dynamics to reduce the incidence of certain STs. Strains of *Campylobacter* are known to vary in their pathogenic potential^23^, with some strains particularly effective at cell invasion^24^. Introducing competitively superior strains into a transmission source presents a way to ensure that particularly pathogenic strains are unable to establish via competitive exclusion, as has been demonstrated in experimental studies^25^. Alternatively, an understanding of these competitive frameworks presents the possibility for the use of live vaccine candidates, whereby bacterial strains that have been weakened can be used to trigger an immune response and limit pathogenic strains^26^. While promising results in such vaccine candidates have begun to appear^27^, reliable effectiveness of these approaches requires knowledge of the underlying population dynamics. As of yet, such dynamics are not properly understood^28^.

Understanding of these mechanisms is further exacerbated due to the fact that the exact route of entry into the flock is still uncertain. While it is generally considered that horizontal transmission is the most likely source of flock infections^29^, with STs carrying over from other locations on a farm, there still exists evidence of some infections caused due to vertical transmission^30^ and wild bird crossover^31^. The possibility of multiple points of entry for *Campylobacter* to enter a flock would explain the inability for improved biosecurity alone to reduce outbreak incidence, and may even suggest that stopping colonisation outright may be a fruitless endeavour, further supporting the need to utilise the manipulation of competitive hierarchies within the host microbiome; if the bacteria cannot be kept out of the farms, perhaps it can yet be kept out of the birds.

Investigations into the varying prevalence of specific STs have shown, experimentally^32^ and numerically^33^, that a multitude of STs can be isolated from a chicken at any given time, and yet within this pattern of co-occurrence only one specific ST will usually be seen to dominate the gut, being isolated in far greater proportions than its co-colonisers. Through this mechanism, a diverse mix of STs can exist in this way within a flock of many chickens, each carrying their own cohort of STs, and each with their own resident dominant strain. This observation constitutes a metacommunity^34^ of STs. A metacommunity is defined as a system where small communities interact with one-another, and influencing the dynamics within each individual community. In our instances, the competing STs within a single host chicken can be thought of as a community, with multiple STs competing for dominance within one chicken, yet the dynamics within each individual chicken influence neighbouring chickens, resulting in a level of flock-wide dynamics as well. By utilising various mathematical frameworks from the wider ecological literature, we can begin to uncover how STs can co-exist within the flock, and to then ascertain what dynamic properties cause some newly introduced STs to die out, and others to persist.

To investigate this dynamic behaviour, this study utilises two mathematical modeling approaches to query the data from a long-term broiler-breeder flock prevalence study by Colles et al. (2015)^35^, which reports the STs isolated from individual birds within a flock across a year. A competition matrix model, such as that outlined by Ulrich et al. (2014)^36^, is used to estimate a global competition matrix, detailing the competitive outcomes of pairwise competition between STs. This matrix quantifies the likelihood of specific competitive outcomes, namely if some STs will always outcompete some other STs, or whether such competitive outcomes can have unpredictable results. More importantly, they also provide insight into the competitive hierarchy seen within the broiler microbiome, whether that be a highly structured hierarchy, whereby dominant STs will always out-compete lesser-able STs in a gradually decreasing order of competitive advantages, or perhaps instead a system of intransitive competition. Intransitive competition, or ‘rock-paper-scissors’ competition, instead is defined as a system whereby loops are observed in the rank of competitive outcomes, for example if ST A outcompetes ST B, ST B outcompetes ST C, and ST C then outcompetes ST A^37^. We refer to this cyclic relationship as an intransitive triad. In such a system, there can be frequent turnover of competing organisms, as no one entity is necessarily globally superior. Intransitive competition has been shown to have far-reaching implications for ecological stability and biodiversity, enabling species coexistence^38^, promoting biodiversity^39^, and enabling species cooperation^40^.

Building on this, we then use the estimated competition matrix within a discrete-time patch-occupancy model to simulate and explore the broader dynamics of how STs move between birds in a flock, displace one another, and capitalise on the niches presented by uncolonised birds. Patch-occupancy models simplify a system to a series of ‘patches’, be it spatial units or, in our case, individual chickens, where each patch can be occupied by only one organism at a time, in our case, the dominant ST of *Campylobacter*. The turnover in occupation by different organisms is captured by a series of probabilistic transition mechanisms, which have had great success in demonstrating persistence within metacommunities^41^, due to minimising the assumptions placed upon the population dynamics of the system. The mechanisms that allow for sustained biodiversity in metapopulation models have been shown to primarily be the demographic factors of transmission and mortality of competing species^42,43^. i.e. how well a bacteria can invade a host, and how well it can remain there. In our case, we consider transmission as a measure of how many subsequent chickens will likely be challenged by the established ST in a host bird in the following timestep, the outcome of such a challenge is then decided by the previously estimated competition matrix. Bacterial mortality meanwhile is considered as the probability that a dominant ST will die out in the subsequent timestep, leaving the host bird susceptible to a new invading ST (not to be confused with bird mortality). By building a simulation of the system from which the data was gathered, we estimate these two specific parameters for each ST, and examine how these vary between STs and how they correlate with the observed frequency of each ST.

By presenting quantified estimates into the growth, spread, and competitive ability of each individual ST, we are able to provide insight into how STs of *Campylobacter* interact with one another, both within a host chicken, and within a flock as a whole.

## Methods

### Data

In the original study, a flock of 500 broiler breeders was monitored, with 200 birds labelled with leg-rings and monitored for a total of 51 weeks. Each week, cloacal swabs were taken from a random selection of 75 of the labeled birds, and tested for the presence of *Campylobacter* through standard culture methods. Positive samples were then genotyped (MLST), enabling the ST and species of the *Campylobacter* isolate to be specified. Note that, while multiple STs can occupy a host-bird simultaneously, it is frequently observed, experimentally^32^ and theoretically^33^, that a single ST will broadly dominate the gut at any given time. Hence the sole ST recorded from a positive bird is a reflection of which STs are most dominantly expressed at that timepoint. Furthermore, these dominant STs in a host bird will dominate for roughly a week before being replaced by a competitor^33^. 39 distinct STs of varying prevalence were observed across the year within the flock, 25 of *Campylobacter jejuni* and 14 of *Campylobacter coli*. 19 of these STs appear very rarely, with less than ten total appearances in the data. Due to this limited number of data, meaningful conclusions as to their competitive abilities cannot be given, and as such we do not consider these STs in our analysis, considering only the 20 STs for which more than ten instances of occurrence were recorded in the data. An example layout of a small portion of this data is presented in Figure 1, and the total prevalence of STs over time is displayed in Figure 2. Negative samples are not shown in Figure 2, as this data is not used for the competition matrix model. Further experimental details can be found in the original publication^35^.

**Figure 1.**
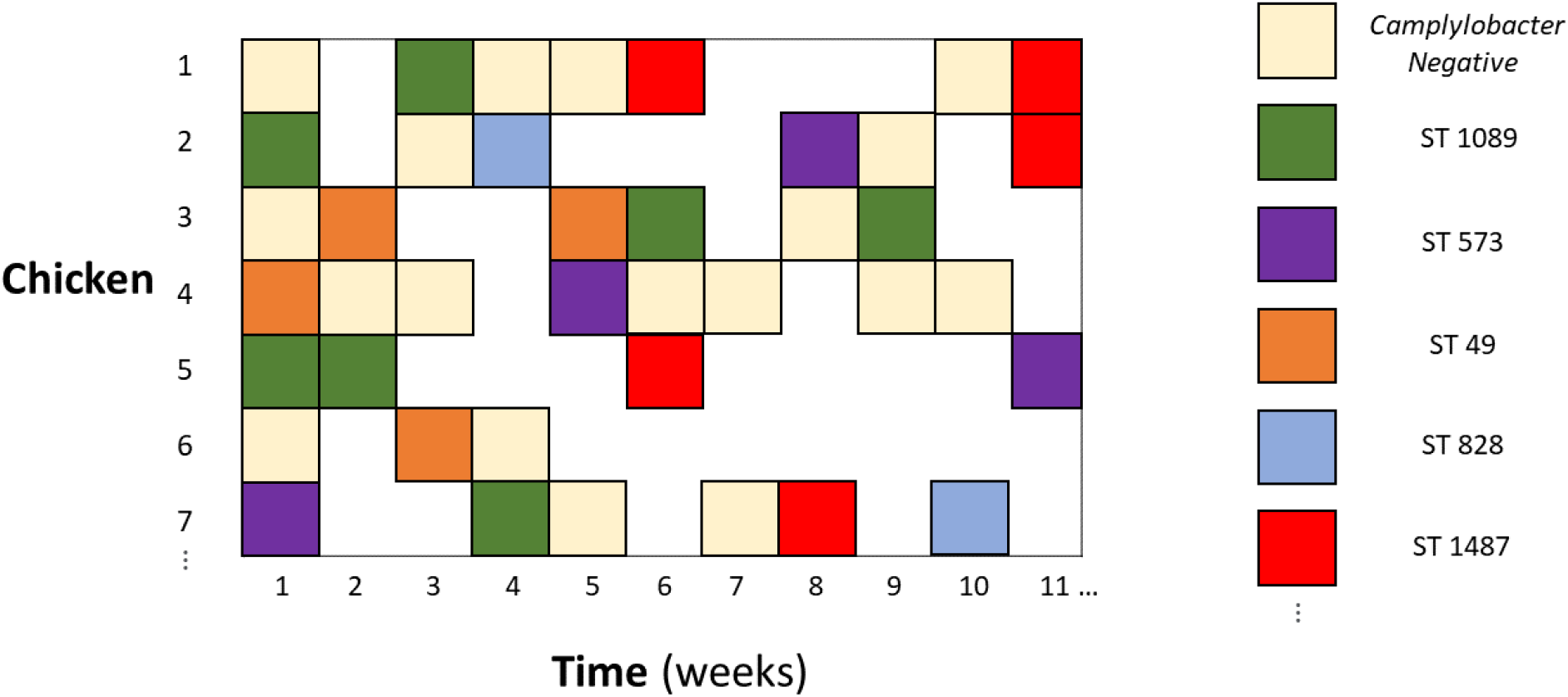
Example portion of the ST prevalence data. From a total flock of 500 broiler breeders, 200 were labelled with leg-rings. These 200 are captured in the rows of the data frame. Each week 75 of these birds were tested for the presence of *Campylobacter* for 51 weeks (columns). Birds were marked as either free from *Campylobacter* (marked in tan), or if found to be *Campylobacter* positive, the sequence type (ST) of the bacteria was recorded. Blank white spaces indicate where a bird was not tested for that particular week. The whole data set comprises 200 rows, 51 columns, and captures 39 distinct STs.

**Figure 2.**
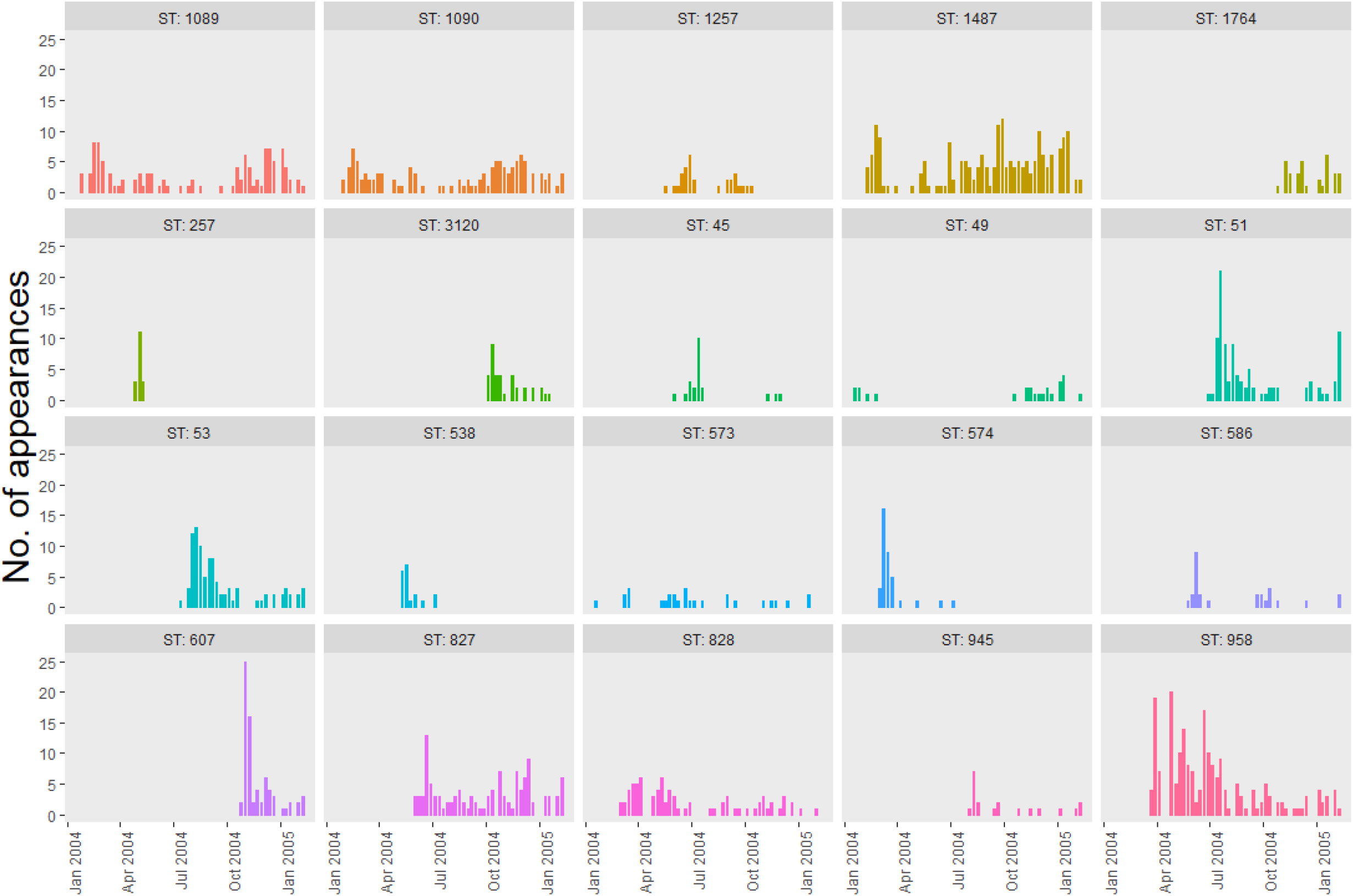
Histograms of ST prevalence across the study period reported in Colles et al. (2015)35. STs with less than 10 appearances are not displayed.

### Competition Matrix

We first estimate a competition matrix, detailing all pairwise competition outcomes between all STs. Formally, we define that, for a system of *n* STs, the competition matrix, *C*, is an *n× n* square matrix where element *C_i, j_* represents the probability that ST *i* out-competes ST *j* in a pairwise competition. By definition, the diagonal elements of *C* are equal to 1, and *C_i, j_* = 1 *−C_j,i_*.

By using the time-series abundance data of all STs throughout the flock, as shown in Figure 2, one may back-infer the pairwise competitive strengths between all STs within the flock. Based upon the methods outlined by Ulrich et al. (2014)^36^, this competition matrix may be estimated by first inferring a transition matrix, *P*: an *n × n* square matrix where *P_i, j_* represents the probability that a chicken colonised by ST *i* is instead colonised by ST *j* in the next time period. Note that this matrix *P* is not the same as the competition matrix *C*, as the observed transitions could represent the result of multiple sequential competitions between STs - the replacing ST has not only outcompeted the present occupant, but also all other incoming STs.

To estimate this transition matrix, *P*, consider an *n ×* 51 frequency matrix *A*, where element *A_i,t_* denotes the number of chickens that ST *i* was isolated from at time *t*, and where *n* is the number of distinct STs. This matrix is directly built from our data, where element *A_i,t_* can be seen as the ‘No. of appearances’ of ST *i* in week *t* from Figure 2. This frequency matrix is then related to our transition matrix, *P*, via the equation;

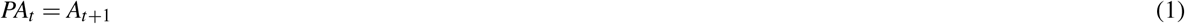

where *A_t_* is a column vector of *n* elements, reporting the abundance of all STs at time *t*. This provides a method by which to estimate *P* by choosing the matrix *P* that best fits equation (1). Homogeneous mixing of STs is assumed, however, another assumption is made in equation (1) that all STs are present and are capable of appearing at each time point. This is not representative of biological reality. We see from Figure 2 that some STs do not appear in the flock until later in the experiment, and while it could plausibly be being out-competed in every prior instance, it is more plausible that the ST has simply not yet infected the flock. As such, we adapt equation (1) by also implementing a binary-filled ‘presence’ matrix *Z*, an *n×* 51 matrix, where element *Z_i, j_* is either 0 or 1, denoting whether or not ST *i* is present in the flock at time *j*. i.e. when a ST is not observed within a flock in a particular week, we do not consider it’s impact on that week’s transition dynamics.

If ST *i* is isolated in the data at time *t*, we mark it as present in matrix *Z* for times *t* through to *t* + 3, to account for the possibility of a ST being reduced to low levels, not captured in the data. This three week window was determined from our earlier numerical simulations33, showing the average duration for which a low-level “non-dominant" ST might persist within a host. We rewrite equation (1) as:

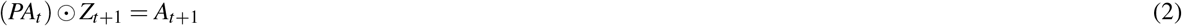

where *Z_t_*_+1_ is the (*t* + 1)^th^ column of *Z*, and ⊙ is the Hadamard (element-wise) product. In essence, *Z* simply acts as a switching mechanism, to switch off the possibility of transitions to a ST that has not yet emerged. This approach carries multiple benefits. Primarily, the transition matrix now represents the transition probabilities for a flock where all STs are present simultaneously. This inference allows more of the dataset to be utilised, without having to divide our experimental data into multiple regions of different sized matrix calculations. A possible limitation to this approach is that it allows inference of competitive outcomes between STs that do not appear at the same time in the original dataset. i.e. it can infer based on the growth abilities of a ST at a later time how it would fare against a ST from an earlier time. While this inference is useful, these limited instances are not experimentally verifiable. As such, we do not display these few “assumed" competitive strengths in our results, to avoid confusion.

Once the best fitting *P* to equation (2) has been found, we may use this *P* to estimate the associated competition matrix *C*. Ulrich et al. (2014)^36^ presents such a methodology whereby, assuming homogeneous mixing, the transition matrix *P* and the competition matrix *C* are linked by the relationship:

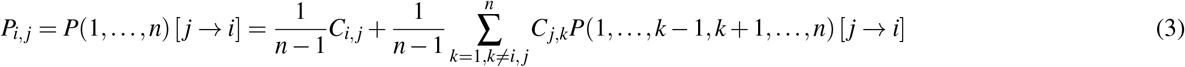

for *i* ≠ *j*, and

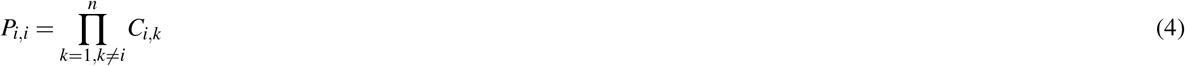

where the range of summation in (4) is calculated across the subset considered in the notation *P*(1*, …, n*). Heuristically, one considers the transition probabilities as the proportional outcomes of all possible competitive interactions. In a four-species system, equations (3) and (4) would define:

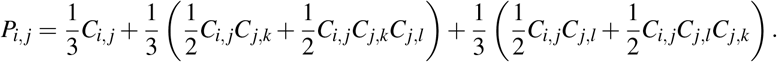

In small systems, the probability of successful transition for each ST could be directly calculated as the proportional outcome of all possible competitive interactions as given in equations (3) and (4). However, for our system of 20 STs this is computationally impossible, as the size of equation (3) will rapidly balloon for such a large system. Instead we therefore used the approximation approach of Ulrich et al. (2014)^36^:

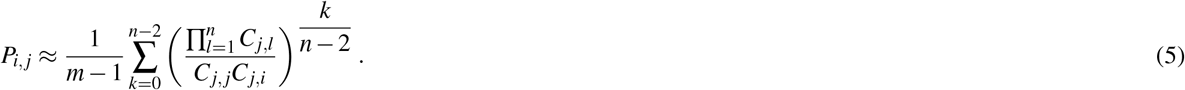

This approximation was found to estimate a randomly drawn 20 *×*20 test competition matrix with a mean value error < 0.001.

The above methodology allows us to choose a trial competition matrix, *C*, convert this to a transition matrix, *P* via equation (5), and then evaluate how well this transition matrix simulates the observed data, *A*, via equation (2). All that is required now is an approach by which to find the “best” competition matrix *C*. As such, we estimate the competition matrix *C* using the above equations within a Bayesian framework, using the Just Another Gibbs Sampler (JAGS) program^44^, a Markov chain Monte Carlo (MCMC) sampling program utilising Gibbs sampling. Specifically the model was called and analysed within R by using the rjags package^45^. We considered wide, uninformative, uniform priors on the elements of *C*. Convergence was considered well-achieved, with every element of *C*’s posterior distribution displaying a potential scale reduction factor (PSRF) < 1.03, and a Monte Carlo standard error (MCSE) less than 5% of the standard deviation of the sample. The code used is made available at https://osf.io/3rd4e/.

Lastly, we quantify the amount of intransitivity observed from the best-fit competition matrix *C*. While many metrics of measuring intransitivity have been proposed^46^, the most suitable is generally considered to be Kendall and Babington Smith’s *d_s_*^47^; a measure of the proportion of three-species intransitive loops found within the competition matrix. i.e. we measure the number of cyclical intransitive triads seen in the competition matrix, and divide this by the total number of possible triads for a competition matrix of that size.

### Patch-occupancy model

The estimated competition matrix gives insight into the interactions between different *Campylobacter* STs, however it cannot by itself answer our questions as to how biodiversity of STs is maintained within the flock. The previous metacommunity modelling studies of May & Nowak (1994)^42^ and Hanksi & Gyllenberg (1997)^43^ have demonstrated that persistence can be largely managed by differences between the colonising ability and mortality of competing organisms. As such, we estimate parameters describing the colonising ability and mortality for each of our 20 considered STs. Figure 2 shows that some STs occur with increased frequency compared to other present STs. For example, STs 1487 and 573 both seem to persist within the flock throughout the entire recorded experimental duration, and yet ST 573 is observed in far fewer birds throughout this time. We hypothesise that differences in the demographic parameters between these STs may explain the differences in the underlying population dynamics.

A patch-occupancy model was designed to simulate the experimental data as closely as possible. In this instance, the patches considered are the 500 chickens that make up the flock, and the STs of *Campylobacter* present are the occupying entities.

A 500 *×* 51 matrix is initialised, where each row denotes a specific chicken in a flock, and each column a time-point (a week), so as to replicate the data structure shown in Figure 1. Element (*i, t*) thus records which ST, if any, has colonised chicken *i* at time *t*. The first column is initialised to match the proportion of STs recorded in the first week of the dataset in Figure 2. Each time-step is then simulated in turn, to iteratively generate the subsequent 50 columns. In each timestep, each established ST may be removed for the following timestep with probability, *μ_i_*, the ST-specific mortality parameter that we seek to estimate. STs that persist to the next timestep then have the opportunity to infect other chickens. The number of other chickens that are challenged by this resident ST is drawn from a Poisson distribution, Pois(*λ_i_*), where *λ_i_* is a ST-specific parameter. Borrowing from the parlance of the ecological literature, we refer to this parameter as the average ‘propagules released’ by ST *i*. If a challenged chicken is currently uncolonised by *Campylobacter*, they then become colonised by the invading ST. If a challenged chicken is currently colonised by a different ST, this is treated as a competitive event, whereby the winner of the pairwise competition will be the occupying ST for the following timestep, and the loser is removed. This outcome is decided by the probabilities estimated in our previous model, given by the matrix *C*.

When new STs appeared for the first time in the experimental data, they are directly introduced into the patch-occupancy model at the proportion and time-step they were first observed. One exception is made for ST 49, which was unobserved for so long in the experimental data, that two specific introduction events were allowed. Appendix 1 outlines the pseudo-code detailing this model structure. The model was programmed in R and the code is available at https://osf.io/3rd4e/.

Considering transition events on the weekly timescale provided in the original data is considered valid based upon theoretical modelling work showing that dominant STs in a host bird will dominate for roughly one week before being replaced by a competitor^33^. Much like our previous model, this provides a framework whereby a trial solution of *μ_i_* and *λ_i_* for each ST *i* can be used, and the resulting ST population dynamics can be compared against the population dynamics observed in the original data. We wish to find the values of *μ_i_* and *λ_i_* that best capture the patterns seen in Figure 2. We score a trial solution by comparing the relative proportions of ST frequency at each time-step with the proportions shown in the original data. The specific iterative framework as outlined in Appendix 1 cannot be integrated into a Bayesian system, so we instead utilise machine learning techniques to seek the optimum solution.

We first find an estimate for the average parameter values across all STs to use as an initial trial solution for each individual ST. We collapse the data to a binary state of either *Campylobacter*-positive or *Campylobacter*-negative, and use simulated annealing to find the average *μ* and *λ* values that best simulate the data, using a scoring function defined by the absolute difference between the infection proportions in every column and every row between the model data and experimental data. This is so that the algorithm selects the parameters that also capture the frequency with which chickens may transition from being *Campylobacter*-positive to *Campylobacter*-negative. This provided a best-fit solution of *μ* = 0.7, and *λ* = 3.2. These values were then used as initialisation points for each ST-specific parameter set (*μ_i_, λ_i_*), which are then iteratively adapted using genetic algorithm approaches to find the best-fit solution.

Genetic algorithms, so named for their inspiration by natural selection, generate “mutations” of the initial trial solutions, and the resulting mutations which best describe the data will in turn inform the next generation of trial solutions. A genetic algorithm of population size 200 was run for 100 iterations, using the (0.7, 3.2) estimate as a suggested population element for each specific ST.

## Results

Figure 3 shows the pair-wise competition values for all STs. STs that do not naturally co-occur during the experiment have been represented with a grey-box, as meaningful conclusions as to their competitive interactions cannot be drawn. The matrix has been re-ordered to maximise the number of values >0.5 in the upper-diagonal, thus showing the identified hierarchy.

**Figure 3.**
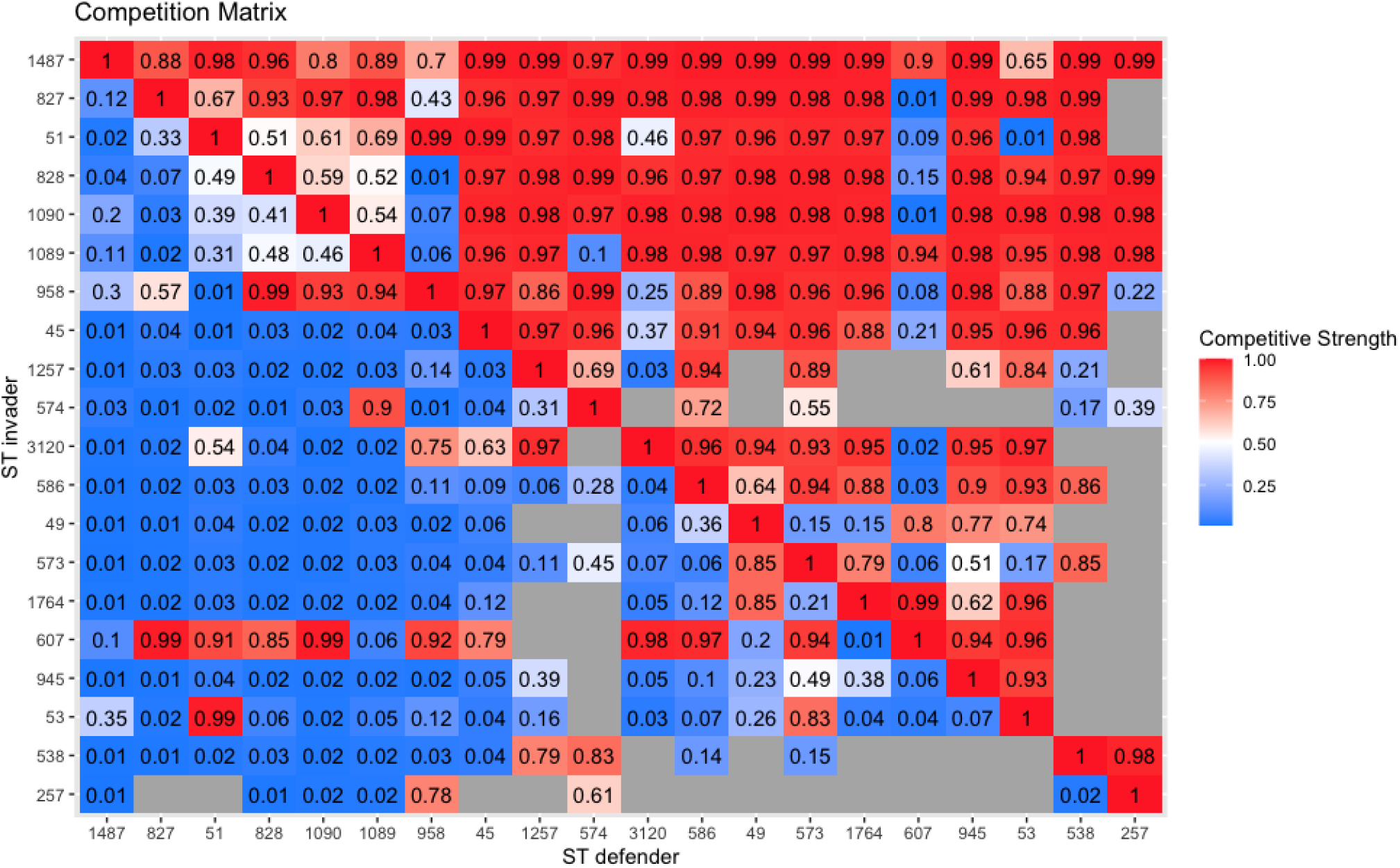
Matrix of pairwise competition strengths between *Campylobacter* STs. Element (*i*, *j*) depicts the probability that ST *i* out-competes ST *j* in a pairwise competition. Empty grey boxes depict cases where two STs do not coexist during the experiment, thus their competitive relationship cannot be estimated. Rows are ordered to maximise the number of values >0.5 above the diagonal. The structure reveals a strong competitive hierarchy, with the strongest competitors at the top of the matrix.

A strong hierarchical structure can be observed, with STs at the top of the matrix mostly outcompeting all STs below them. Some intransitive loops can be seen within the matrix however, for example ST 607, which is able to out-compete some STs higher up the hierarchy. When uniformly sampling the missing values of the matrix shown in Figure 3, an average of 125 intransitive triads are recorded for the competition network, compared to a hypothetical maximum of 330 for a (complete) 20 *×* 20 matrix, resulting in an intransitivity score of *d_s_* = 0.379 (Kendall and Babington Smith’s *d_s_*^47^). In comparison, on sampling 100,000 random 20 *×* 20 competition matrices, the lowest number of intransitive triads generated was 196, hence our observation of only 125 triads supports a system of significant hierarchical competition.

The competition matrix shown in Figure 3 is then utilised within the patch-occupancy model to estimate ST-specific transmission and mortality parameters. These parameters are displayed below in Figure 4. Mortality (*μ*) we define as the probability that an established ST will die-out from its host bird naturally from one week to the next. To capture ST-specific transmission effects we report the average propagules released (*λ*), the average number of other chickens that an occupying ST will challenge for the following timestep, with the outcome of these challenges decided by the above competition matrix.

**Figure 4.**
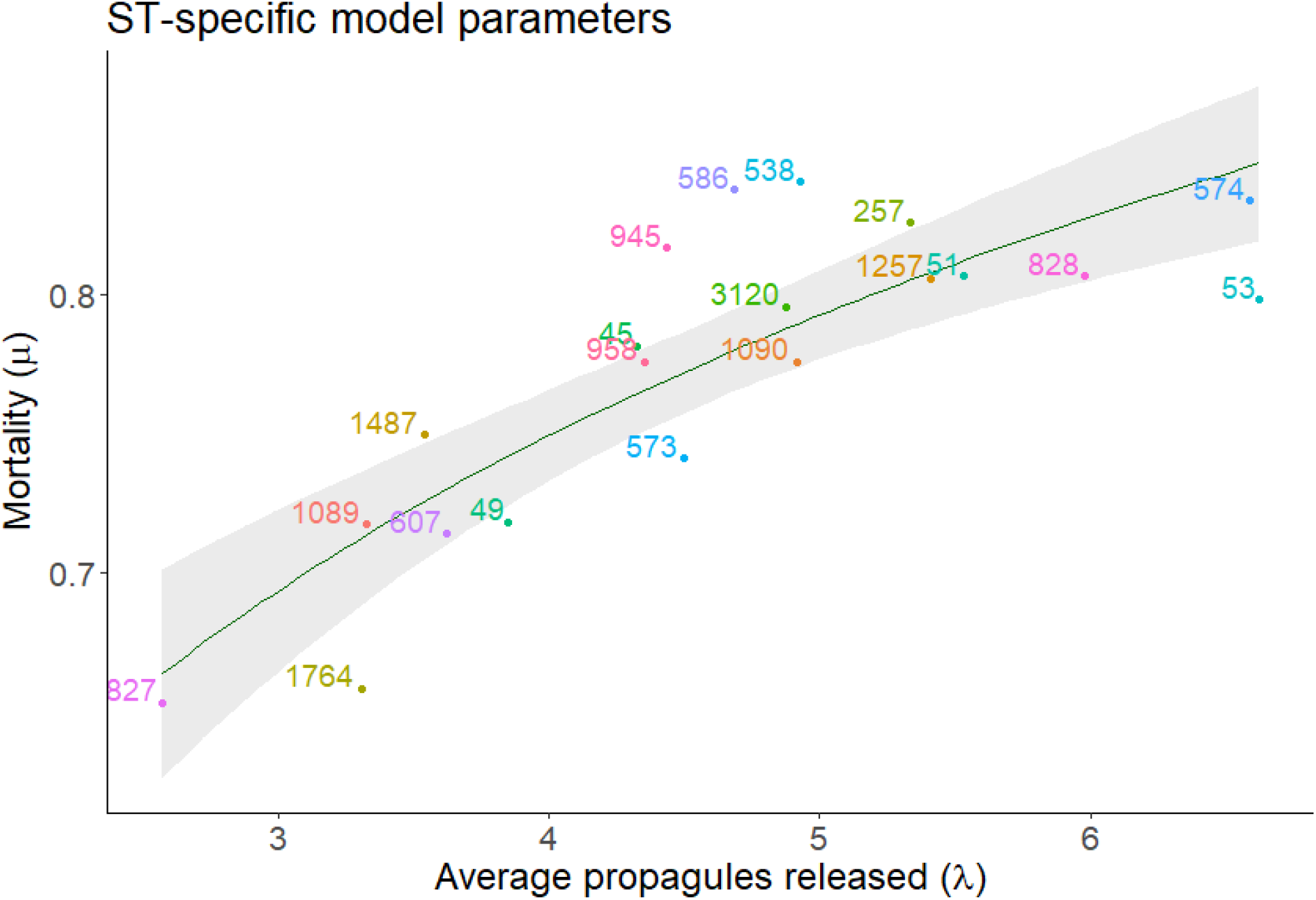
ST-specific model parameters for patch-occupancy model. Mortality (*μ*) depicts the probability that a ST dies out from one time-point (a week) to the next. If the ST does not vacate a host, it releases propagules that challenge other host chickens. The average number of chickens challenged (*λ*) is a model parameter depicted on the *x*-axis. The green line displays the statistically significant (*p* < 0.0001) logarithmic regression between the two variables. We see a positive trend whereby higher mortality is compensated by a greater number of propagules being released.

The positive logarithmic trend (*p <* 0.0001) shows a relationship whereby STs with a higher mortality (they die out more frequently) can maintain their presence in the flock by being able to colonise more chickens.

## Discussion

Here we have investigated the ecological drivers maintaining *Campylobacter* diversity within chicken flocks. By quantifying competition, transmission, and mortality parameters through two mathematical frameworks, we have highlighted the demographic differences between *Campylobacter* sequence types, and shown that the metacommunity of STs operates within a strict competitive hierarchy, with some STs capable of outcompeting other STs, and hence replacing them as the dominant strain within host birds.

The competition matrix shown in Figure 3 effectively disproves the hypothesis that ST diversity may have been maintained by a system of intransitive competition, as very few intransitive triads were found within the system. Intransitive loops have been shown to theoretically support coexistence of many competing organisms, dependent on growth rate differences and intransitive cycle length^48^, and such effects have been demonstrated in small plant communities^49^. Despite the wealth of theoretical work surrounding the impact of intransitive competition, real-world evidence of such systems is lacking. An experimental study searching for such effects across five different taxonomic groups by Soliveres et al. (2018)^50^ was unable to find strong evidence of intransitivity in any of their studies other than experiments with mosses, and found zero intransitive triads in their bacterial experiment. As such, our inability to identify clear signs of intransitivity is unsurprising. Only STs 53, 607, and 3120 showed clear evidence of being able to out-compete STs higher in the hierarchy. All three of these STs appear to have remained prevalent in the flock from their point of entry to the end of the experiment time, possibly suggesting that STs that are able to form an intransitive loop may be more capable of invading and persisting in the flock.

Within this competitive hierarchy, we also show that the magnitude of the respective competitive probabilities are relatively large. In the the upper diagonal of Figure 3 values are greater than 0.9, suggesting that the competitively superior STs not only outcompete a vast number of other STs, but that they outcompete these other STs decisively, winning competitive interactions over 90% of the time in most instances. This is in line with experimental studies into inter-strain competition, with El-Shibiny et al. (2007)^51^ demonstrating how a strain of *Campylobacter* is not only able to outcompete a multitude of other strains, but to do so repeatedly in multiple experiments.

This evidence of a clear competitive hierarchy further stresses how specific mechanisms must underpin the observed maintenance of biodiversity of *Campylobacter* STs. Under such competitive conditions, biodiversity of a metacommunity has been shown to be feasibly maintained by trade-offs between transmission and mortality^42, 52–54^. Under such a system, the co-occurrence of multiple STs can be explained by competitively strong STs displaying high mortality rates, namely that after replacing a resident ST, they naturally die out from the host quickly. Alternatively, their transmission ability may be compromised such that, although they may be very effective competitors, they are unable to proliferate as fast as other STs, and thus may not challenge a high number of other chickens from one week to the next. Likewise, a competitively weak ST, such as ST 53 in Figure 3, may not be able to withstand competition from incoming STs, but is able to persist in the flock by challenging a higher number of chickens each week (high number of propagules released), and surviving within these host birds for a longer period of time (low mortality). The patch-occupancy model presented was designed to specifically quantify these mortality and mean propagule release parameters, and are presented in Figure 4.

Figure 4 shows that all STs can be placed somewhere within a life-history trade-off. In general, STs displaying high mortality, may persist in the environment by releasing a higher number of average propagules, and vice-versa. May & Nowak (1994)^42^ theoretically showed that for a newly emerging entity into a community to successfully invade a metacommunity, and to then persist, they need to fill a yet unrepresented area of this transmission-mortality spectrum. i.e. to persist, they need to have no close neighbours in the plot of Figure 4. This may be demonstrated by STs 827 and 53. Both STs can be seen from Figure 2 to appear within the flock mid-way through the time span, and to then successfully persist through to the end of the experiment. Both of these STs can also be seen from Figure 4 to be outliers on the transmission-mortality spectrum, with ST 827 having the lowest mortality of all observed STs, and ST 53 having the highest number of mean propagules released. As a further interesting contrast, competitive ability does not appear to have influenced this, as ST 53 is one of the weakest competitors in the metacommunity, and ST 827 is one of the strongest, as shown in Figure 3.

However, this mechanism alone has historically been unable to account for the vast amount of sustained biodiversity observed in nature. Building on the theoretical findings of May & Nowak (1994)^42^, Bonsall et al. (2004)^53^ demonstrated that species within a hierarchical competition structure, competing for the same resource, may co-exist by clustering into ‘life-history guilds’. Competitively strong species may simultaneously co-exist by sharing similar demographic parameters. At the same time, competitively weaker species will also persist in the environment, by also sharing similar demographic capabilities with one another. Scheffer and van Nes (2006)^54^ highlighted the same result, concluding that newly emergent species would only persist in the environment if either (i) they were significantly competitively superior to all existing species, or (ii) if they were similar enough to existing species, both competitively and demographically, so as to exist within this particular life-history guild niche. Our results however do not show evidence of such ecological guilds.

Figure 4 shows that, while STs do form a life-history trade-off, STs appear in a broadly even distribution across this mortality-propagule trade-off. Furthermore, some STs that appear to be demographically similar vary greatly in their competitive ability and respective population dynamics. From Figure 2, we can broadly delineate STs by four distinct dynamic profiles: a ST may either persist in a flock or die out, and it may exist at high-frequency or low-frequency. It was assumed that one could characterise these four distinct dynamic profiles by their competition, average propagule release, and mortality parameters, and yet no such pattern has been found in this study. For example, the STs 257, 574, 45, and 1257, could all be characterised as appearing in high frequency, before then dying-out. Yet despite these similar dynamical behaviours, all STs place broadly across the competition-propagule-mortality spectrum, with no common trends in their placement. Likewise, STs 586, 573, and 945 could all be categorised as persisting in the flock, though recovered at low frequency, and yet all three STs are found in broadly different placements in Figures 3 and 4. In general, STs that appear in high frequency appear to correlate with higher competitive potential in Figure 3, though no such trend can be associated with persistence.

Since these STs do not demonstrate the guild-assemblage ‘clumping’ structure in Figure 4 (shown by Bonsall et al. (2004)^53^ to be necessary for biodiversity maintenance in this instance), it suggests that some other mechanism must be enabling the co-occurrence and persistence of *Campylobacter* STs. Based upon the broader wealth of investigations into *Campylobacter* dynamics, we can posit three potential hypotheses driving these clearly seen differences in population dynamics between STs:

i. Host-bird variability. It has been shown in numerous patch-occupancy systems that patch quality (meaning that some patches are ‘easier’ to colonise than others) can have a tremendous impact on the overall population dynamics, having even greater impact than differences between how patches are connected^55, 56^. Yu & Wilson (2001)^57^ theoretically showed that while differences in life-history trade-offs were necessary for co-existence, significant heterogeneity in patch quality or density was necessary to support a large number of species. Such patch variation also made it possible for newly emergent species to persist even if the species was inferior in both competitive and colonisation ability. In our context, variation in patch quality and density would translate to host birds varying in their response to bacterial challenge, with some chickens ‘easier’ to colonise than others. Indeed, through Bayesian transition models we have shown using this same data set in Rawson et al. (2020)^58^ that a flock contains a mixture of birds that are highly resilient to bacterial challenge, and highly susceptible birds that operate as ‘super shedders’. These super shedders are consistently being colonised by a variety of *Campylobacter* STs with high turnover. Poor individual bird health and welfare has been previously shown to correlate with a reduced immune response, with measures such as stocking density^59, 60^, food withdrawal, and heat stress61 all contributing to increased *Campylobacter* colonisation. Yu & Wilson’s (2001)^57^ study directly shows that the host-bird variation seen in Rawson et al. (2020)^58^ removes the need for newly emerging STs to be sufficiently similar to persist in the flock. This further supports the idea that the proliferation of *Campylobacter* in a flock is influenced primarily by the individual birds.
ii. Seasonal variation. Broiler flock colonisation by *Campylobacter* has been well-documented to follow a seasonal trend^62, 63^, with flocks more likely to become colonised in the warmer summer months than the winter. The data behind this modelling study was gathered over 51 weeks, January 2004 to January 2005, so would plausibly have been impacted by seasonal variation. The original study examining the impact of local environmental variables on the data set we have considered^35^ (and subsequent Bayesian transition analyses^58^), were unable to identify any temporal trend within the total *Campylobacter* prevalence, however the *Campylobacter coli* STs did appear less frequently during the summer. It is thus plausible that seasonal variation may have impacted the population dynamics of the occupying STs in the flock via some yet-unidentified mechanism. An example of this may be seen by comparing the population dynamics of STs 53 and 574. Both STs occupy a similar placement in the propagule-mortality spectrum of Figure 4, and yet, despite ST 574 being more competitively able than ST 53, ST 574 does not persist in the flock, while ST 53 does. One possible explanation for this is that ST first appeared within the flock in July, while ST 574 appeared in February.
iii. Stochasticity. While our patch-occupancy model is a probabilistic one, the mechanisms by which a metacommunity of *Campylobacter* STs persist is determined by a number of random events. The events of a ST first entering the flock, chickens ingesting colonised faeces, and of then establishing themselves within the microbiome all encompass a wide number of stochastic events which could change the resulting population dynamics. Coward et al. (2008)^28^ showed that attempts to replicate population dynamics of *Campylobacter* within broilers were largely unsuccessful, even in the most simple cases of just two competing strains. They posited that this was likely due to “founder effects”, small variations in population level at first inoculation which could have large consequences for the flock-wide population dynamics. We have previously shown this effect through a series of stochastic differential equations in Rawson et al. (2019)^33^, whereby a variety of overall population dynamics can be observed dependent on stochastic events when the population of a *Campylobacter* ST is very low. Likewise, upon running the patch-occupancy model for the estimated parameters presented in the results, some STs would persist in some actualisations, but not others. Thus, attempting to characterise some dynamical profiles by mortality and transmission parameters, may not be possible as our experiment displays only one dynamic outcome of many possible ones.

One important caveat to this work must be stressed. Since broiler flocks are slaughtered anywhere from 5 to 11 weeks of age, longitudinal studies into the *Campylobacter* population dynamics are not possible, birds are not alive for long enough for us to observe long-term dynamics from which to extract parameter estimates. As such, this experimental data was gathered from a flock of broiler-breeders, the birds that lay the eggs that become broiler flocks. As we have discussed above, host bird factors may have significant implications for the overarching population dynamics of the microbiome, meaning that these estimated parameters could plausibly be different in commercial broiler flocks. Broiler and breeder flocks are kept under slightly different housing conditions and diet provisions^64^, and breeder flocks have also been shown to shed smaller amounts of *Campylobacter* than commercial broilers^65^. Since this study has focused on investigating *Campylobacter*-specific factors, our conclusions remain relevant to commercial broiler flocks, namely that the population dynamics remain deeply susceptible to impact from a variety of factors, such as season and host bird health.

The primary finding of this work highlights how the life-history trade-offs we have identified fail to provide an explanation for the persistence and co-occurrence of multiple *Campylobacter* STs. This further supports the notion that suppressing and controlling outbreaks of *Campylobacter* cannot be achieved through bio-security alone, and reflects calls for a ‘One Health’^66^ approach, whereby further understanding is needed of how *Campylobacter* and broilers interact and affect each other. We have shown that demographic advantages alone cannot determine which STs of *Campylobacter* will come to dominate a flock of chickens, and that it may instead come down to a ST being in the right place at the right time, or rather, the right chicken in the right season.

## Author contributions statement

F.M.C. collected the data. T.R., J.C.D.T., and M.B.B. conceived the study. T.R. built the models and wrote all associated code. T.R. wrote the manuscript. F.M.C., J.C.D.T., and M.B.B. supervised the project. All authors aided in interpretation of results and reviewed the manuscript.

## A Appendices

## A.1 Appendix 1 Patch-occupancy model pseudo-code

**Algorithm 1:**
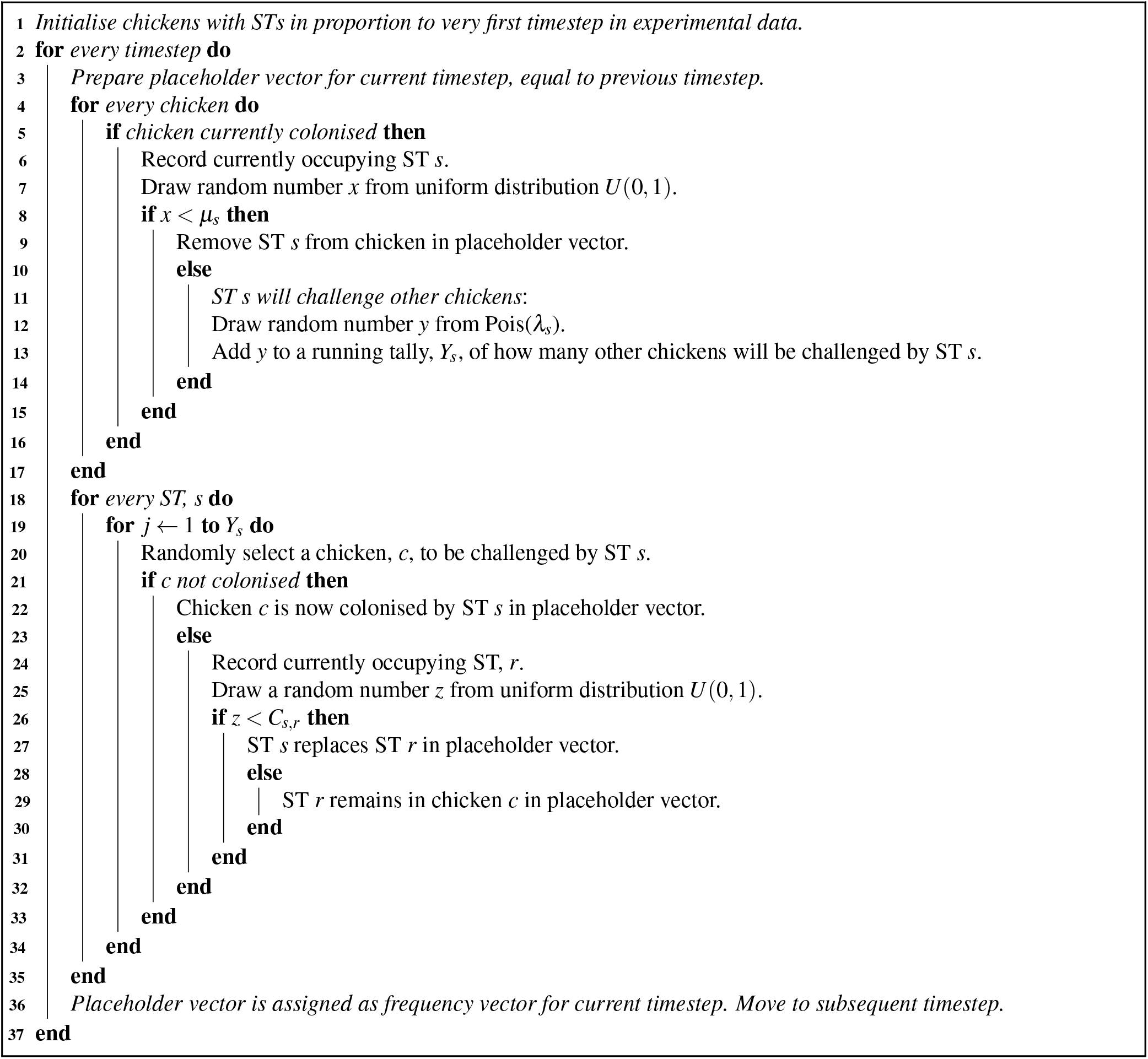
Patch-occupancy model pseudo code

